# Nocturnal intraocular pressure rise is regulated by norepinephrine via RHOB

**DOI:** 10.1101/2025.03.21.644679

**Authors:** Keisuke Ikegami, Takumi Takahashi, Atsuro Oishi, Akihide Yoshimi, Miki Nagase, Atsuya Miki, Shinobu Yasuo, Satoru Masubuchi

## Abstract

Intraocular pressure (IOP), a key factor in glaucoma development, is regulated by aqueous humor (AH) dynamics, with inflow from the ciliary body and outflow through the trabecular meshwork (TM). IOP has a circadian rhythm entrained by sympathetic norepinephrine (NE) from the superior cervical ganglion. Herein, we investigated its underlying regulatory mechanisms in the TM. Through comprehensive gene expression analysis of human TM cells and mouse eyes, we identified 18 genes upregulated by NE stimulation, including the small GTPase RAS homologous protein B (*RHOB*). Promoter assays revealed *RHOB* upregulation via the cyclic adenosine monophosphate (cAMP) response element on its promotor. NE stimulation for 6–9 h increased RHOB level and cellular adhesion, and suppressed liquid permeability in the TM cells, indicating a time-dependent effect. RHOB deficiency increased TM macrophage phagocytosis and eliminated NE-induced suppression of phagocytosis and permeability, whereas RHOB overexpression had the opposite effect. Instillations of RHO or RHO-kinase inhibitors to mice eye reduced nocturnal and NE-induced IOP elevation. Our findings suggest that NE can elevate IOP via RHOB-mediated inhibition of TM phagocytosis, positioning RHOB as a potential glaucoma treatment target and IOP rhythm regulator.

## Introduction

In most organisms, circadian rhythms govern numerous physiological processes. In mammals, the suprachiasmatic nucleus (SCN), functions as the primary circadian pacemaker. It receives optical input from the retina to regulate daily biological rhythms^1^. The SCN coordinates the rhythmic activity of peripheral tissues and cells through intricate pathways involving the autonomic nervous system^2^. Norepinephrine (NE), also known as noradrenaline, is released from the superior cervical ganglion (SCG), a component of the sympathetic nervous system, and conveys circadian signals to the ciliary body of the eye, thereby influencing pupil size and other ocular functions^3^. Moreover, glucocorticoids (GCs) released from the adrenal glands under the control of the SCN via the hypothalamus-pituitary-adrenal (HPA) axis serve as potent endocrine timekeepers. They play a critical role in synchronizing circadian activity, with glucocorticoid receptors widely expressed across various peripheral tissues^4^.

Although glaucoma is a leading cause of blindness in older individuals, it still has no effective cure. Elevated intraocular pressure (IOP) is a key factor in the onset and progression of glaucoma, which is characterized by gradual vision loss. The IOP is regulated by aqueous humor (AH) dynamics and follows a circadian rhythm^5^. In humans, IOP increases at night, regardless of posture^6^. Nocturnal IOP elevation occurs in both diurnal^7^ and nocturnal animals, with the process being controlled by the SCN in mice^8^. Patients with glaucoma also exhibit nocturnal IOP elevation^6,9^ with phase shifts observed in primary open-angle glaucoma (POAG) and normal-tension glaucoma (NTG)^9^. Abnormal IOP rhythms are linked to NTG^10^, and aging desynchronizes IOP rhythms in older individuals^11^. Disruption of this rhythm in night-shift workers is associated with a higher risk of optic nerve damage and glaucoma^11,12^. These findings highlight the need to regulate the nocturnal IOP for effective glaucoma management and underscore the importance of circadian mechanisms governing AH dynamics. Our previous research showed that NE and GCs convey circadian timing signals to the eye, thereby affecting IOP rhythms^13^. However, their role in regulating phagocytosis in the trabecular meshwork (TM), which influences IOP, remain unclear^14,15^. NE is known to inhibit macrophage phagocytosis^16^, and our previous study suggested that impaired AH drainage may partially explain the nocturnal IOP elevation in mice. We found that β1 adrenergic receptor (β1AR)-mediated suppression of TM phagocytosis occurs via non-genomic pathways involving the activation of the cAMP-exchange proteins directly activated by cAMP (EPACs) pathway^17^. However, the genomic mechanisms responsible for the regulation of TM phagocytosis remain unknown (Fig. 1A). Interestingly, although the cAMP-PKA pathway attenuates TM phagocytosis, protein kinase A (PKA) does not appear to influence the SHIP1-PIP3 pathway^17^, suggesting that PKA inhibits TM phagocytosis via gene regulation. Indeed, suppression of TM phagocytosis by NE was partially but dose-dependently restored by the transcriptional inhibitor cordycepin (Fig. 1A, Extended Data Fig. 1).

**Fig. 1:**
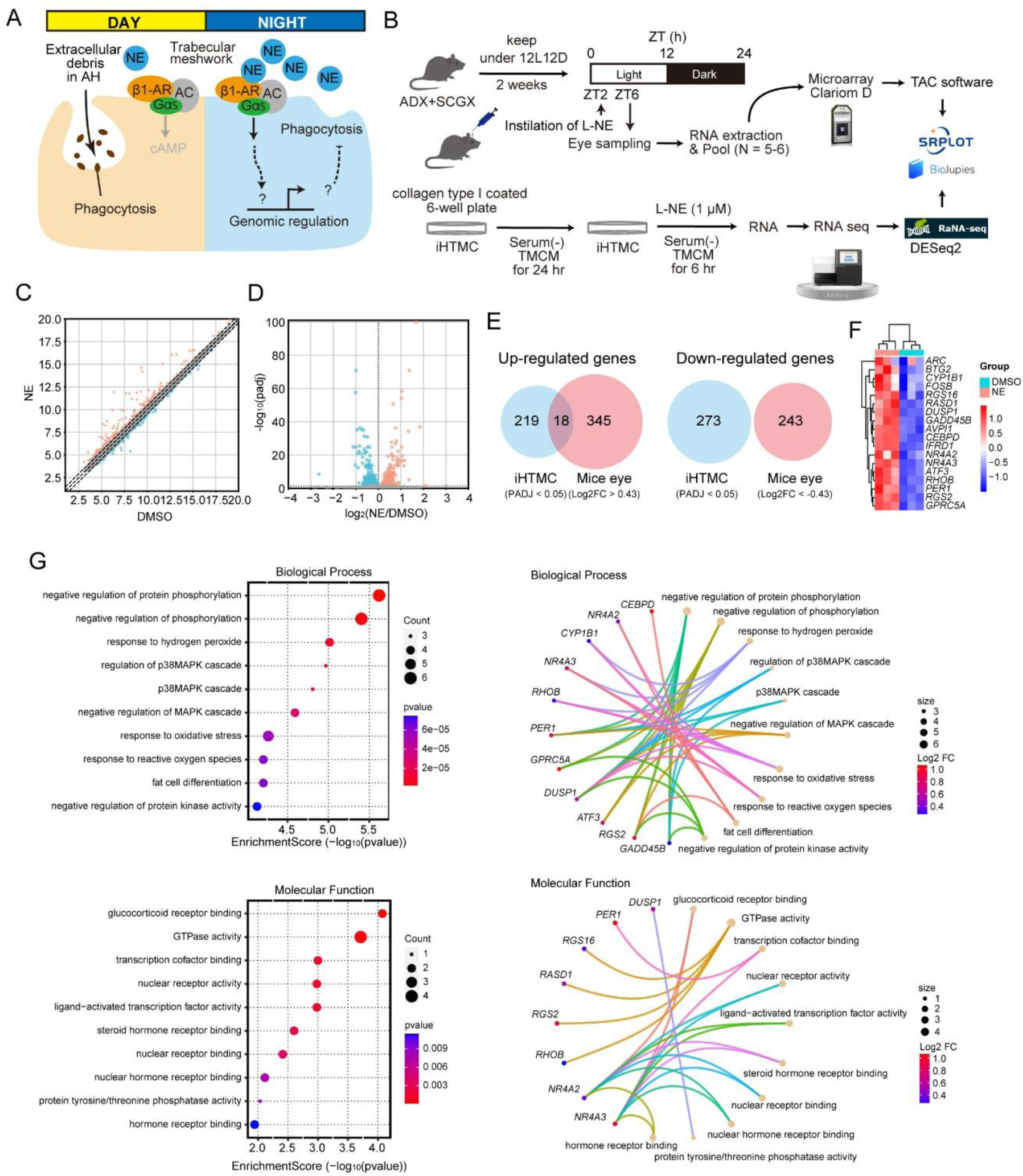
Norepinephrine (NE) stimulation upregulated small GTPase-related genes in mouse eye and human trabecular meshwork (TM) cells. A Nocturnal NE elevates intraocular pressure via macrophage phagocytosis in the TM; however, its genomic regulatory mechanisms remain unknown. B Microarray analysis of NE-treated adrenalectomized (ADX) and superior cervical ganglionectomized male mice, and RNA-seq analysis of NE-treated immortalized human TM cells (iHTMCs). NE was instilled into the mouse eye at ZT2 and RNA was extracted from the eyes at ZT6. ZT, Zeitgeber time. Pooled RNA was used for microarray analysis. C Scatter plot of differentially expressed genes (DEGs) in NE-treated eyes compared with control eyes. The plot shows microarray analysis results with upregulated genes (363) marked in red and downregulated genes (243) marked in green. The DEG threshold was set to |log_2_FoldChange| > 0.43. D Volcano plots of DEGs with three-fold lines (PADJ < 0.05). Upregulated genes (237) are marked in red and downregulated genes (273) are marked in green. E Venn diagrams DEGs revealed 18 commonly upregulated genes. F Cluster heatmap of the 18 genes in iHTMC upregulated by NE. G Gene ontology (GO) enrichment analysis of the 18 DEGs. The top ten (ranked by p-value) significantly enriched GO terms in the biological process and molecular function groups for each gene are shown. Pearson’s correlation matrix of GO enrichment analysis of DEGs. GTPase activity is associated with multiple genes. Larger symbols indicate a greater number of genes correlated with the GO term.

The AH dynamics are regulated by the AH inflow^18^ and outflow^19,20^. The outflow pathway is responsible for the homeostatic regulation of IOP and is mainly regulated by the coordinated generation of AH outflow resistance mediated by the constituent cells of the TM and Schlemm’s canal (SC)^19,20^. Phagocytosis in TM macrophages can decrease particulate material and debris from the AH, attenuate outflow resistance, and contribute to IOP reduction^19^. Phagocytosis is thought to play an important role in the normal functioning of the outflow pathway by keeping drainage channels free of debris. Conversely, actin remodeling such as fragmentation or polymerization regulates AH outflow, and the actin cytoskeleton of TM cells is a therapeutic target in patients with glaucoma. Small GTPase RAS homologous protein (RHO)-associated coiled-coil-containing protein kinase (ROCK) inhibitors lower the IOP by relaxation of the TM via disruption of actin stress fibers and by activation of phagocytosis^21–23^. However, because all phagocytic processes are driven by a finely controlled rearrangement of the actin cytoskeleton^24,25^, the separate effects of these two processes are difficult to verify. In addition, changes in TM cell characteristics, such as increased cell adhesion, contractility, and stiffness, can lead to a higher resistance to AH outflow^26^, which results in elevated IOP.

In addition, NE suppresses wound macrophage phagocytic efficiency through α- and β-AR-dependent pathways^16^, and can promote actin polymerization in the retina^27^. NE has been shown to increase the adhesion of several cells^28^. In the TM, cAMP/PKA activation and downstream RhoA inactivation lead to the loss of actin stress fibers, focal adhesions, and disassembly of the matrix network^29^. However, the detailed effects of NE on TM function have not been elucidated. Hence, this study aimed to uncover the regulatory mechanisms of NE in TM cellular functions that regulate nocturnal IOP. We explored the key genes involved in NE-mediated AH outflow in diurnal IOP changes in mice and immortalized human TM cells, and identified their regulatory pathways. Gene modification and pharmacological approaches were used to identify NE-mediated TM cellular physiological functions. Furthermore, the roles of the identified molecules in IOP rhythm regulation were assessed by pharmacological instillation in mice.

## Results

### NE upregulates several genes in the TM

To determine the effects of NE stimulation on gene expression in the mouse eye and human TM cells, we performed a comprehensive gene expression analysis on NE-treated mice and immortalized human TM cells (Fig. 1B). Differential gene expression analysis was performed using microarray analysis in the mouse eye (PRJDB20328; |Log2FC| < 0.43; Fig. 1C) 4 h after administration of NE eye drops, which is the period when the IOP increases after NE exposure^13^, and RNA sequencing (RNA-seq) of human TM cells (PRJDB20330; PADJ < 0.05; Fig. 1D) 6 h after NE exposure. We did not detect any common downregulated genes; however, 18 common upregulated genes (*GPRC5A, RGS2, PER1, RHOB, ATF3, NR4A3, NR4A2, IFRD1, CEBPD, AVPI1, GADD45B, DUSP1, RASD1, RGS16, FOSB, CYP1B1, BTG2,* and *ARC*) were detected (Fig. 1E, F). Gene ontology (GO) enrichment analysis indicated negative regulation of phosphorylation, response to hydrogen peroxide, regulation of p38MAPK cascade, and negative regulation of MAPK cascade as enriched biological processes; glucocorticoid receptor binding, GTPase activity, and transcription cofactor binding were among the enriched molecular functions. Glucocorticoid receptor binding (*NR4A2* and *NR4A3*) may support the NE/Dex interaction, in which Dex and NE suppress phagocytosis in a dose-dependent manner, consistent with previous *in situ* studies^30,31^. We focused on small Rho-associated GTPase activity (*RGS16, RASD1, RGS2*, and *RHOB*) related to the ROCK pathway (Fig. 1G). Ras superfamily GTPases are molecular switches that cycle between GDP-bound inactive state and GTP-bound active state to control many signaling pathways. Furthermore, Kyoto Encyclopedia of Genes and Genomes (KEGG) pathway enrichment analysis strongly suggested that the amphetamine addiction and circadian entrainment pathways (Extended Data Fig. 2A), as well 18 genes, were associated with ganglion- and eye-related cells (ciliary or mast cells) by enrichment analysis of Tabula Sapiens and Jensen TISSUES (Extended Data Fig. 2B,C). Thus, it is plausible that, though RHOB activity like most Rho GTPases is regulated by GTP/GDP loading, NE-induced increase in GTPase activity may be involved in the nocturnal increase in IOP observed in mice.

### NE increases RHOB expression in the TM

To identify the signal transduction pathways related to the NE-upregulated genes, we performed enrichment analysis of kinase perturbations. We found significant similarity between the 18 upregulated genes and the group of genes whose expression is decreased by Rho-associated protein kinase 1 (*ROCK1*) knockdown (GSE34769) (Fig. 2A) and those whose expression is increased by *ROCK2* knockdown (Fig. 2B). This implies that the ROCK1 pathway is more similar to the genes altered by NE than to ROCK2, and ROCK1 can regulate NE-induced 18 genes. Pathway enrichment analysis of the Reactome using 18 genes revealed several RHO-related indices (Fig. 2C). Since ROCK inhibitors decrease IOP in mice^13^ and reanalysis of previous single-cell RNA-seq data suggests similar *RHOB* expression patterns in almost all human TM cells (Extended Data Fig. 3)^32^, and *RHOA* and *RHOC* were not upregulated by NE (Fig. 1), we focused on RHOB. Indeed, when we measured RHOB activity in NE-treated immortalized human TM cells (iHTMCs), we detected an increase in its activity (Fig. 2D). Western blot analysis of NE-treated mouse eyes showed that NE treatment induced an increase in RhoB protein levels in the whole mouse eye (Fig. 2E). Since the TM involved in IOP control is a very narrow and small region, it is unclear whether NE actually affects RHOB expression in TM cells; therefore, iHTMC were used to verify this. RHOB expression gradually increased after NE stimulation, peaking 6 h after stimulation (two-way ANOVA [interaction, *p* = 0.0419; time, *p* = 0.0597; DMSO vs. NE, *p* = 0.0007]; Fig. 2C,D). Treatment with the β1AR agonist dobutamine for 6 h also enhanced RHOB expression in the TM (Extended Data Fig. 4). These results suggest that NE increases protein level-dependent RHOB activity in the TM.

**Fig. 2:**
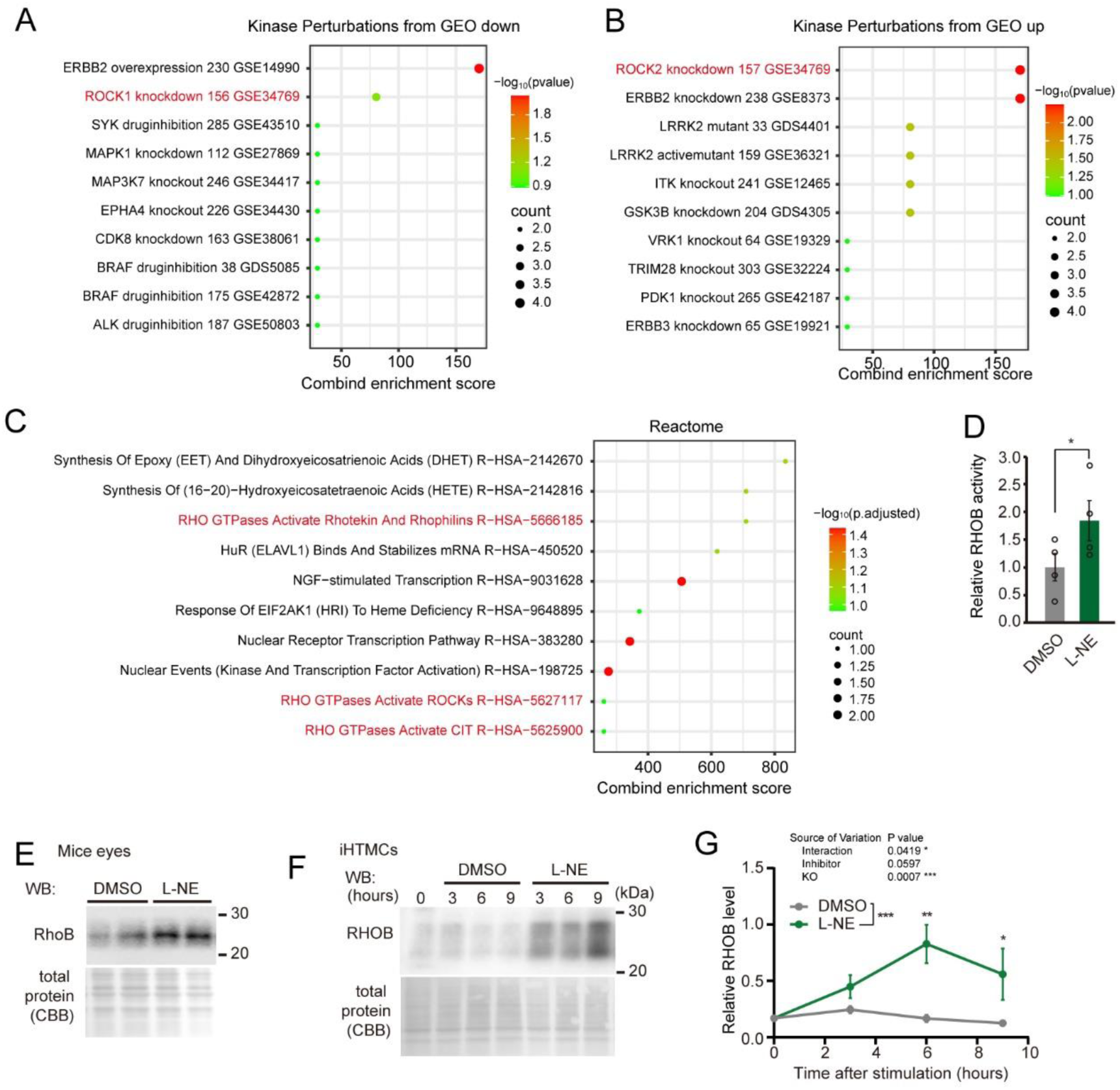
Norepinephrine (NE) treatment induced RHOB expression in mouse eyes and immortalized human trabecular meshwork cells (iHTMCs). **A,B** Top 10 signatures from kinase perturbations extracted from GEO down and GEO up, respectively, linked by the combined score with p-values and gene counts. **C** Pathway enrichment analysis of the Reactome using 18 genes revealed several RHO-related indices **D** RHOB GTPase activity assay of NE-treated iHTMCs showed that NE enhanced RHOB activity (unpaired *t*-test, * *p* < 0.05). Data are presented as bar graphs (mean ± SEM) with scatter plots of independent experiments (n = 4). **E** RhoB protein expression in mouse eyes exposed to L-NE for 5 h. Coomassie Brilliant Blue staining was performed to confirm total protein levels. **F** RHOB induction in iHTMC 3, 6, and 9 h after L-NE administration. **G** Relative RHOB levels by NE exposure time. RHOB expression gradually increased with NE exposure, peaking at 6 h after stimulation (\**p* < 0.05 vs. WT, Šidák multiple comparison test, two-way ANOVA [interaction, \**p* = 0.0419; time, *p* = 0.0597; DMSO vs. NE, \*\*\**p* = 0.0007]).

### NE upregulates *RHOB* expression via CRE

*RHOB* expression is known to be regulated by ultraviolet irradiation, ERK, AKT, and Rho; however, the regulatory pathways related to ARs remain unknown^33,34^. To investigate the regulatory mechanism of *RHOB* expression in the TM, we first performed transcriptional factor (TF) enrichment analysis of ENCODE and ChEA consensus TFs using ChIP-X (Fig. 3A). The cyclic adenosine monophosphate (cAMP)-responsive element (CRE)-binding protein CREB1 (ChEA) was identified as the most important TF for NE-mediated *RHOB* expression (Fig. 3A), indicating the involvement of CREB or cAMP in *RHOB* regulation. In fact, β1AR is a typical G protein-coupled receptor (GPCR) that preferentially binds to stimulatory G protein G_s_ to induce cAMP production. cAMP binds to and activates PKA or exchange proteins directly activated by cAMP (EPACs)^35^. β1AR agonists (L-NE and dobutamine) stimulate the downstream activation of CREB in the iHTMC^17^. In addition, intracellular cAMP accumulate through the action of cAMP inducers forskolin and β1AR agonists^17^.

**Fig. 3:**
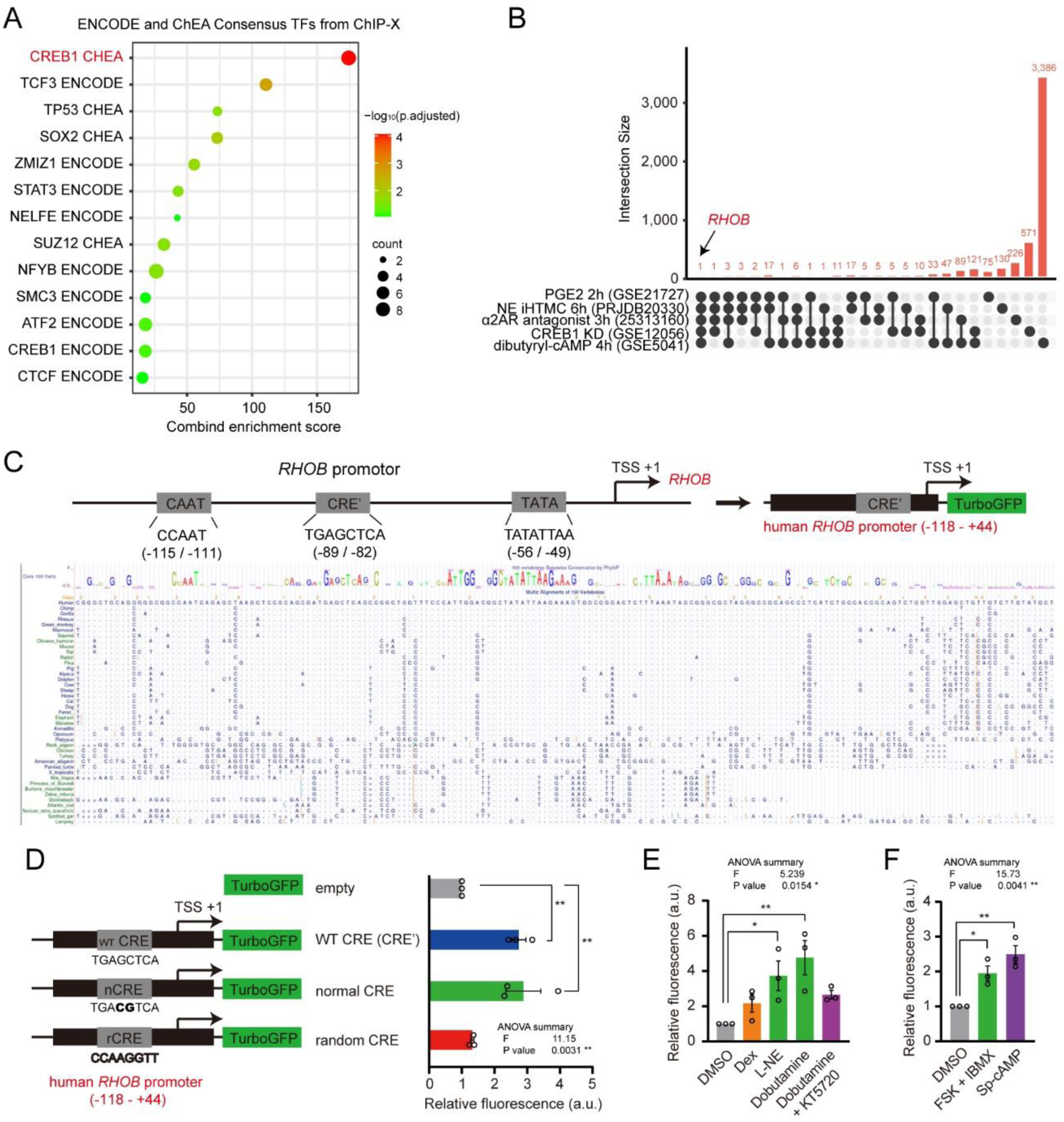
Norepinephrine (NE) upregulates RHOB in immortalized human trabecular meshwork cells (iHTMCs) through the cAMP-PKA-CREB pathway. **A** Transcriptional factor (TF) enrichment analysis of ENCODE and ChEA Consensus TFs from ChIP-X. The cyclic adenosine monophosphate (cAMP)-responsive element (CRE)- binding protein CREB1 has been identified as the most important TF. **B** Upset plot of the cAMP-related five gene sets obtained from GSE21727, GSE12056, 25313160, GSE5041, and PRJDB20330. Only RHOB was identified as the common variable gene. Black points indicate intersections and gray points indicate no intersections. **C** Annotation analysis of the human RHOB promoter (−1,500 to +1 bp) using the UCSC genome browser. The TATA box, CAAT element, and a previously unknown CRE′ site (TGAGCTCA) near the transcription start site (TSS). **D** Mutation promoter assay for CREB-binding sites in the hRHOB gene (−118 to +44 bp). Plasmids without any promoter or enhancer were used as negative controls. Plasmids were constructed with wild-type (WT) CRE′, normal CRE (TGACGTCA) by reversing the middle GC, and random CRE (CCAAGGTT). TurboGFP expression was monitored in iHTMCs after NE stimulation. Dobutamine exposure (1 μM) for 4 h induced GFP expression (\**p* < 0.05, \*\*\**p* < 0.001 vs. DMSO, one-way ANOVA [\*\**p* < 0.001], Dunnett’s multiple comparison), while random CRE had no effect (*p* > 0.05). Values represent the means ± SEM of three independent experiments performed in triplicate. **E** NE and dobutamine induced RHOB expression (\**p* < 0.05, vs. DMSO, Dunnett’s multiple comparison test), but dexamethasone (Dex) did not. **F** Sp-cAMP and forskolin [FSK] with 3-isobutyl-1-methylxanthine [IBMX] also induced RHOB expression (\**p* < 0.05, vs. DMSO, Dunnett’s multiple comparison test). Data are presented as scatter plots with mean ± SEM (n = 3 independent experiments).

To explore the importance of the CREB-*RHOB* pathway, we analyzed cAMP- or CREB-related transcriptome data from The Signaling Pathways Project (http://www.signalingpathways.org/ominer/query.jsf) in addition to our data from NE-treated human TM cells. These included 226 upregulated genes in iHTMC after 6 h of NE exposure [PRJDB20330] (Fig. 3B), 75 upregulated genes in human primary osteoblasts after 2 h of exposure to prostaglandin E2 (PGE2)-activating Gs-coupled GPCRs (EP2 or EP4 receptor) (GSE21727; FDR < 0.05, |Log2FC| > 2), 130 upregulated genes in K562 human leukemia cells by CREB1 knockdown (GSE12056; PADJ < 0.05, |Log2FC| > 2), 571 upregulated genes in rat liver cells 3 h after injection of Gi-coupled α2AR blocker chlorpromazine-increasing cAMP (25313160; PDR < 0.05, |Log2FC| > 2), and 3,386 upregulated genes in mouse immortalized preadipocytes 4 h after dibutyryl-cAMP exposure (GSE5041; FDR < 0.05, |Log2FC| > 1). Among these gene sets, only RHOB was detected as a common gene (Fig. 3B), suggesting an important relationship between RHOB regulation and the cAMP-CREB signaling pathway.

The presence of CRE, which bind CREB or CREB family members^36^, has not been reported on the RHOB promoter in any animal. Annotation analysis of the human *RHOB* promotor (−1,500 bp to +1 bp) using the UCSC genome browser revealed a previously unknown CRE′ site (TGAGCTCA; −89 bp to −82 bp) near the transcription start site that may be no longer capable of binding CREB^37^, which is highly conserved in vertebrates, especially in mammals (Fig. 3C). The *RHOB* promoter only contains this CRE′ site. To directly investigate the role of CRE in the NE-mediated increase in *RHOB* promoter activity (−118 bp to +44 bp), we constructed a series of plasmids containing reporters with wild-type (WT) CRE′, normal CRE (TGACGTCA) by reversing the middle GC and random CRE (CCAAGGTT), and then monitored TurboGFP expression after NE stimulation in iHTMC (Fig. 3C,D). As expected, GFP expression was induced in iHTMC transfected with plasmids containing either the WT or normal CRE *RHOB* promotor, 4 h after dobutamine (1 μM) treatment (*p* < 0.01, vs. empty, Dunnett’s multiple comparison test, Fig. 3D), whereas random CRE had no effect on NE-induced GFP expression (*p* > 0.05, Fig. 3D), suggesting the involvement of CRE in NE-induced *RHOB* expression. Using the WT CRE promotor, NE and dobutamine induced *RHOB* expression (*p* < 0.05, vs. DMSO, Dunnett’s multiple comparison test, Fig. 3E), but the glucocorticoid receptor agonist dexamethasone (Dex), an IOP rhythm regulator^13^, did not (*p* > 0.05, vs. DMSO, Fig. 3E). Furthermore, KT5720, a PKA inhibitor, suppressed dobutamine-induced *RHOB* CRE activity (Fig. 3E). Sp-cAMP (a cAMP analog) and cAMP inducers (forskolin [FSK] with 3-isobutyl-1-methylxanthine [IBMX]) also induced RHOB expression (*p* < 0.05, vs. DMSO, Dunnett’s multiple comparison test, Fig. 3F), indicating the direct effect of cAMP on *RHOB* expression. Taken together, these findings suggest that NE can induce *RHOB* expression via the cAMP-CREB-CRE′ site in mammals.

### NE attenuates permeability in the TM

The TM forms a layered structure, and AH is expelled through gaps between the layers^38^. To evaluate AH permeability under conditions that mimic the *in vivo* TM, we cultured TM cells in layers using culture inserts and constructed an evaluation system for cell permeability by measuring the difference in static pressure caused by the color of the medium and the height of the liquid surface (Fig. 4A). Using this system, we examined the effect of dobutamine on liquid permeability in the TM layer and its time dependency. Permeability decreased gradually with the duration of dobutamine exposure, with the maximum permeability reduction at 6 h, then recovering upon longer exposure times (Fig. 4B). These results suggested that NE may prevent AH outflow by increasing cell adhesion and cell-to-cell adherence.

**Fig. 4:**
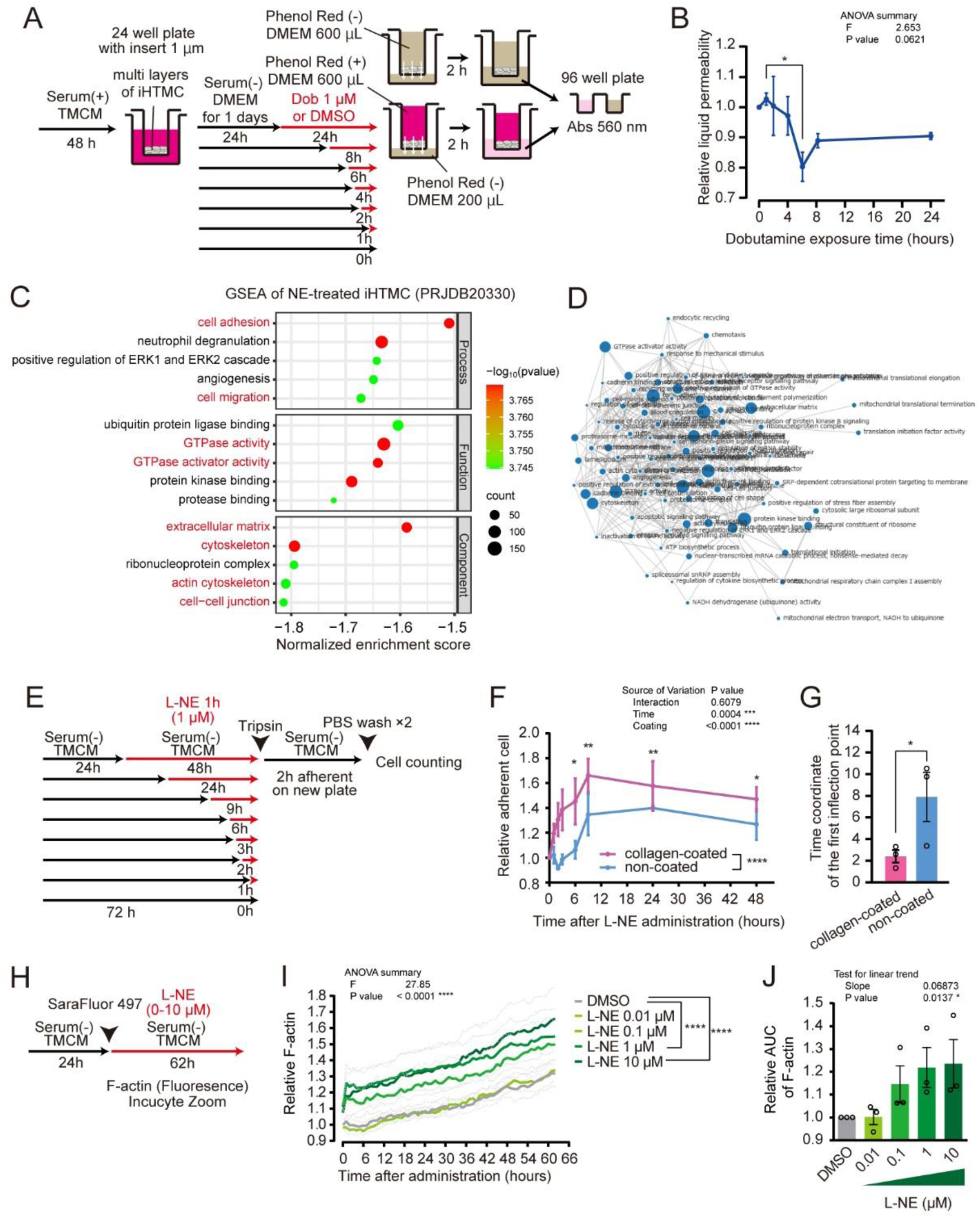
Norepinephrine (NE) inhibits permeability in multilayered immortalized human trabecular meshwork cells (iHTMCs). **A** Experimental scheme to verify liquid permeability in multilayer iHTMC culture using culture inserts. **B** Temporal changes in liquid permeability of dobutamine-treated iHTMC by exposure time. Minimum permeability was observed after 6 h of dobutamine stimulation (\**p* < 0.05 vs. 6 h, Dunnett’s multiple comparison test). Data are presented as line graphs (mean ± SEM) from independent experiments (n = 3). **C** Gene set enrichment analysis of NE-treated iHTMC. **D** Network diagram of the gene ontology terms. Circle size indicates the number of genes. **E** Schematic of the cell adhesion assay to investigate the effect of L-NE exposure time on iHTMC. **F** Temporal profile of adherent cells. NE treatment increased the number of adherent cells, which peaked at 9 h and then gradually decreased (\**p* < 0.05, \*\**p* < 0.01, vs. 0 h, Dunnett’s multiple comparison test, two-way ANOVA [interaction, *p* = 0.6079; time, \*\*\**p* = 0.0004; coating, \*\*\*\**p* < 0.0001]). Data are presented as line graphs (mean ± SEM) from independent experiments (n = 3). **G** Time coordinates of the first inflection point after logistic growth fitting. NE treatment promoted the increase in cell adhesion (\**p* < 0.05, unpaired t-test). **H** Schematic of the actin polymerization analysis using an F-actin probe in iHTMC. **I** Temporal changes in F-actin fluorescence in iHTMC. The immediate increase in actin polymerization after NE stimulation was dose dependent (\*\*\*\**p* < 0.0001 vs. DMSO, one-way ANOVA [\*\*\*\**p* < 0.0001], Dunnett’s multiple comparison). **J** Area under the curve of the F-actin change. NE dose-dependently increased actin polymerization (\**p* < 0.05, linear trend test, one-way ANOVA [\**p* = 0.0137]). Data are presented as bar graphs (mean ± SEM) with scatter plots of data from independent experiments (n = 3).

Because terms related to cell adhesion and cell-to-cell adherence were not identified using GO analysis (Fig. 1G), we interpreted enrichment results using gene set enrichment analysis (GSEA). GSEA revealed biological processes, molecular functions, and cellular components related to GTPase activity, GTPase activator activity, cell adhesion, cell migration, extracellular matrix, cytoskeleton, actin cytoskeleton, and cell-cell junctions (Fig. 4C,D, Extended Data Fig. 5). This supports previous findings that NE may modulate cell adhesion and actin polymerization in the TM^28,39^.

In our previous report, we showed that transient NE stimulation temporarily suppressed the iHTMC phagocytosis, with a trough at 9 h after stimulation^17^. However, in humans, *RHOB* is expressed not only in macrophages but also in several types of TM cells, such as endothelial cells of Schlemm’s canal (Extended Data Fig. 3)^32^. Thus, *RHOB* may also affect the cytoskeleton and cell adhesion in the TM. We evaluated cell–cell adhesion and the ability of cells to adhere to the extracellular matrix and found that NE stimulation increased the number of adherent cells in a time-dependent manner, peaking at 9 h and then gradually decreasing (Fig. 4E,F). Transient NE stimulation also increased the number of adherent cells (Extended Data Fig. 6). The adhesion-promoting effect was reduced in the absence of an extracellular matrix in a non-coated dish (Fig. 4F,G). These results demonstrate that extracellular matrix-cell adhesion and cell-to-cell adhesion increase after NE stimulation. Furthermore, when we measured temporal changes in actin polymerization in iHTMC using an F-actin probe (Fig. 4H), we observed an immediate dose-dependent increase in actin polymerization after NE stimulation which did not show circadian rhythms (Fig. 4I,J). Taken together, these results indicate that NE inhibits phagocytosis, cell adhesion, and actin polymerization in TM cells in a time-dependent manner.

### RHOB affects TM cell viability

To understand the role of RHOB in TM cellular physiological functions, we first generated *RHOB* knockout (KO) iHTMC by transfection of Cas9-expressed CRISPR plasmid and puromycin-mediated selection (*p* > 0.05, Fig. 5A). RHOB was deleted in iHTMCs (Fig. 5B), and RHOB KO showed no effect on cell growth (Fig. 5C), indicating cell growth mechanisms and the cell cycle in TM cells is independent of RHOB, unlike previous reports on other cells^40^. Furthermore, dobutamine inhibited iHTMC growth, whereas Dex accelerated it (two-way ANOVA [KO; *p* < 0.0001]), which was consistent with previous reports^41,42^. RHOB KO had no significant effect on cell growth, although it slightly enhanced the effect of Dex (*p* < 0.05 vs. WT, Šidák multiple comparison test, Extended Data Fig. 7A,B). In contrast, dobutamine exposure gradually decreased cell viability in a dose-dependent manner in a serum-free medium (Fig. 5D) as previously reported^17^, and RHOB KO slightly rescued this effect (Fig. 5D), indicating the importance of RHOB in NE-suppressed TM cell viability.

**Fig. 5:**
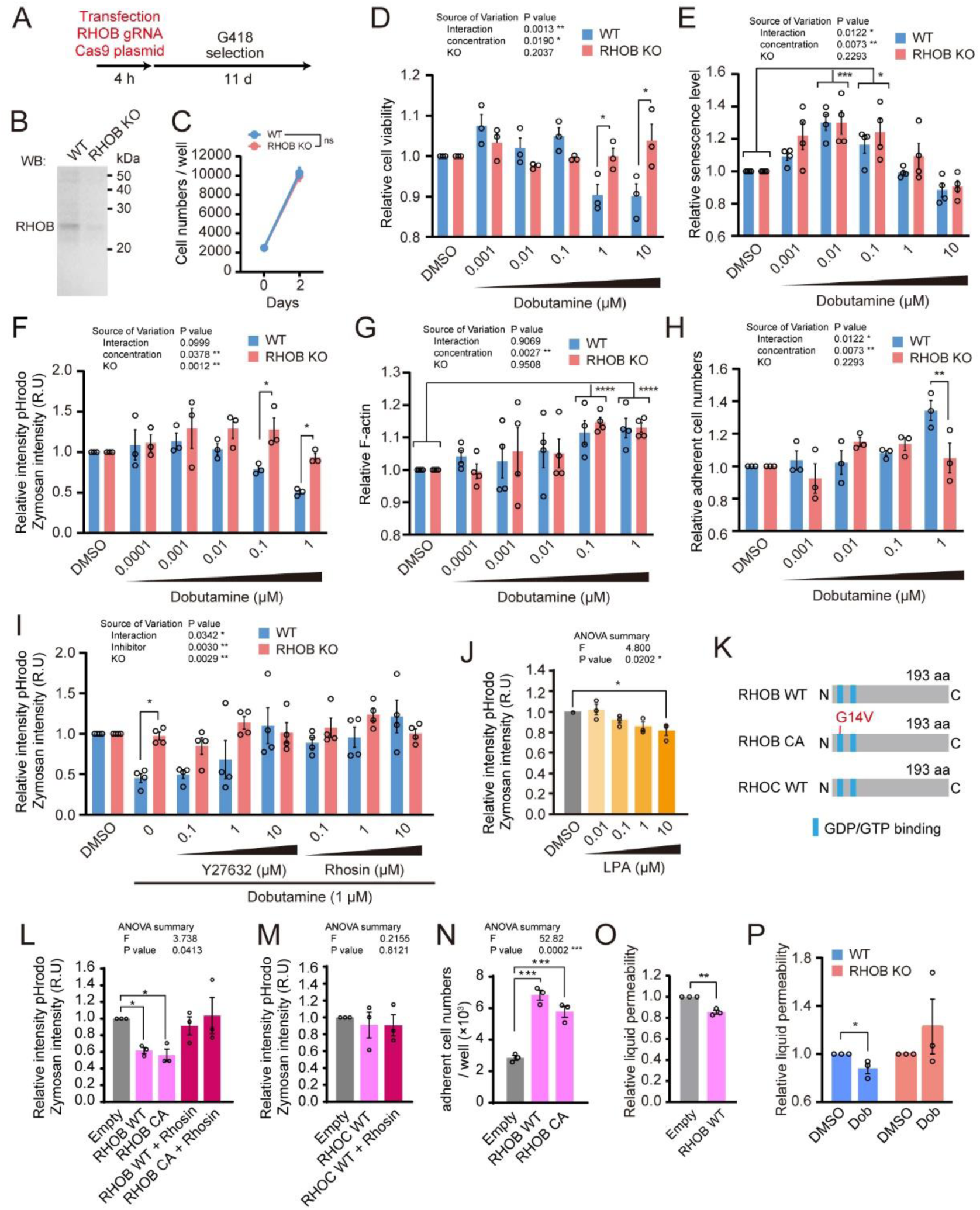
β1AR mainly attenuated phagocytosis through upregulating RHOB. **A** RHOB knockout (KO) was achieved by transfection of CRISPR/Cas9 plasmid. **B** Western blot analysis of WT RHOB and RHOB-KO in iiHTMCs. **C** Cell growth of RHOB-KO iHTMC. KO had no effect on cell growth (*p* > 0.05). **D** Effect of dobutamine on RHOB KO cell viability. Incubation with dobutamine for 6 h dose-dependently decreased cell viability (two-way ANOVA [concentration; \**p* = 0.0190]), whereas RHOB KO showed no effect. **E** Effect of 2 d of dobutamine treatment (0–10 μM) on TM aging. Low concentrations of dobutamine (10–100 nM) increased iHTMC senescence (* *p* < 0.05, *** *p* < 0.001, vs. DMSO, Dunnett’s multiple comparison test, two-way ANOVA [interaction, \**p* = 0.0122; time, \*\**p* = 0.0073; coating, *p* < 0.2293]). **F** Effect of dobutamine on TM phagocytosis in RHOB KO cells. RHOB KO rescued dobutamine-suppressed TM phagocytosis (\**p* < 0.05, vs. WT, Šidák multiple comparison test, two-way ANOVA [concentration; \**p* = 0.0378, KO; \*\**p* < 0.0012]). **G** Dobutamine gradually increased TM F-actin levels, but RHOB KO showed no effect (\*\*\*\**p* < 0.0001 vs. DMSO, Dunnett’s multiple comparison test, two-way ANOVA [concentration, \*\**p* = 0.0027; KO, *p* = 0.9508]). **H** Effect of dobutamine on TM aging in RHOB-KO cells. KO showed slightly decreased TM senescence at high dobutamine levels (\*\**p* < 0.01 vs. DMSO, Dunnett’s multiple comparison test, two-way ANOVA [interaction, \**p* = 0.0122; concentration; \*\**p* = 0.0073; KO, *p* = 0.2293]). Data were normalized using signals from DMSO-treated cells and presented as bar graphs (mean ± SEM) with scatter plot of independent experiments (n = 3–4). **I** ROCK inhibitor Y27632 and RHO inhibitor rhosin rescued dobutamine-attenuated TM phagocytosis (\**p* < 0.05, vs. WT, Šidák multiple comparison test, two-way ANOVA [interaction, \**p* = 0.0342; concentration, \*\**p* = 0.0030; KO, \*\**p* = 0.0029]). **J** Lysophosphatidic acid (LPA), a RHO activator, dose-dependently attenuated TM phagocytosis (\**p* < 0.05 vs. DMSO, Dunnett’s multiple comparison test, one-way ANOVA [\**p* = 0.0202]). **K** Constitutively active (CA) RHOB (G14V in the GDP/GTP binding site) and RHOC WT structures. **L** Overexpression of RHOB WT and CA significantly inhibited phagocytosis, which was rescued by rhosin (\**p* < 0.05, vs. DMSO, Dunnett’s multiple comparison test, one-way ANOVA [\**p* = 0.0413]). Overexpression of WT RHOC had no effect on phagocytosis. **N** RHOB overexpression dramatically increased the number of adherent cells (\*\*\**p* < 0.001 vs. DMSO, Dunnett’s multiple comparison test, one-way ANOVA [\*\**p* = 0.0002]). **O** RHOB overexpression inhibits liquid permeability in iHTMC multilayers (\*\**p* < 0.01, unpaired t-test). **P** Dobutamine-attenuated TM permeability was not observed in RHOB KO iHTMC multilayers (\**p* < 0.05, unpaired *t*-test).

Long-term exposure to Dex can induce TM senescence^43^ and fibrillation^44,45^. Our results showed that iHTMC senescence was slightly increased by Dex in a dose-dependent manner (two-way ANOVA [*p* = 0.0271]) (Extended Data Fig. 7C). Although β-AR activation induces cellular senescence^46^, the effect of NE on TM aging remains unclear. When iHTMCs were incubated with dobutamine (0–10 μM) for 2 d, only low concentrations of dobutamine (10–100 nM) increased senescence, and RHOB KO had no effect on this increase (Fig. 5E). Although we still do not know the time-dependent effect of dobutamine exposure, these results indicate that RHOB KO has no effect on cell growth or NE-induced TM aging.

### β1AR decreases TM phagocytosis via RHOB

Macrophage phagocytosis, cell adhesion, and remodeling of the actin cytoskeleton in TM cells are related to AH outflow^19,21–23^. When we investigated the effect of RHOB KO on these cellular functions, only the phagocytotic activity in TM cells was enhanced by RHOB KO (*p* < 0.05, Extended Data Fig. 8A), but not actin polymerization (Extended Data Fig. 8B) and the number of adherent cells (Extended Data Fig. 8C). Dobutamine dose-dependently decreased TM phagocytosis and increased actin polymerization and TM cell adhesion (Fig. 5F,G,H). However, these effects were significantly inhibited by RHOB KO (Fig. 5F,G,H), and a pharmacological approach using inhibitors for RHO (rhosin) and ROCK (Y27632) rescued the dobutamine-attenuated TM phagocytosis in a dose-dependent manner (Fig. 5I), suggesting the importance of RHOB in TM phagocytosis. Dynasore, an inhibitor of dynamin GTPase activity, has been widely studied in clathrin-mediated endocytosis and phagocytosis in HTMC^17,47^. RHOB KO attenuated dynasore-suppressed phagocytosis in iHTMC (Extended Data Fig. 8D). Dynamin interacts with the actin filaments when the edges of the phagocytic cup close^48^. RHOB does not appear to alter the phagocytic cup but modulates phagocytic cup formation.

In addition, since activation of GPCR β1AR accumulates cAMP in the iHTMC^17^, cAMP accumulation by cAMP inducers (forskolin with IBMX) dose-dependently decreased TM phagocytosis in WT iHTMC, while in RHOB KO iHTMC, low cAMP accumulation dramatically increased TM phagocytosis and high cAMP accumulation arrested dobutamine-suppressed TM phagocytosis (Extended Data Fig. 8E). Furthermore, cAMP accumulation increased actin polymerization, but not TM cell adhesion, unlike the effect of dobutamine stimulation (Extended Data Fig. 8F,G). RHOB KO reduced cell adhesion in a dose-dependent manner, but not actin polymerization (Extended Data Fig. 8F,G), indicating that β1AR modulates TM phagocytosis and cell adhesion via the cAMP-RHOB pathway. In fact, blocking of cAMP prevents β2-AR- and PGE2-suppressed neutrophil phagocytosis^49^, while NE suppresses TM phagocytosis via PKA activation^17^.

### RHOB overexpression decreases TM phagocytosis

In contrast, lysophosphatidic acid (LPA), an activator of RHO protein, dose-dependently attenuated TM phagocytosis (*p* < 0.05, one-way ANOVA; Fig. 5J). To determine the effect of RHOB overexpression on TM functions, we constructed a series of plasmids expressing WT RHOB or constitutively active (CA) RHOB by substituting glycine-14 with valine (G14V). The RHOC construct, which has a high homology of 85%^50^, was used as the control (Fig. 5K). RHOB WT overexpression slightly increased TM cell growth, while RHOB CA and RHOC showed no effect (Extended Data Fig. 9A,B). The bioluminescence signal from the Nano-luc marker showed significant expression of these protein (Extended Data Fig. 9C), we have monitored phagocytosis and cell adhesion in transfected cells (Fig. 5L–5O). Overexpression of RHOB WT and CA decreased TM phagocytosis, as expected (*p* < 0.05 vs. empty, Dunnett’s multiple comparison test), which was rescued by rhosin administration (Fig. 5L), but not that of RHOC (Fig. 5M). The cell adhesion assay revealed that the overexpression of RHOB WT and CA increased the number of adherent TM cells (*p* < 0.001 vs. empty, Dunnett’s multiple comparison test; Fig. 5N). In addition, overexpression of RHOB WT and CA significantly enhanced actin polymerization (*p* < 0.05, *p* < 0.01 vs. empty, Dunnett’s multiple comparison test; Extended Data Fig. 9D), although there was no significant difference in actin polymerization in RHOB-KO iHTMC (Extended Data Fig. 8B). Thus, NE-induced RHOB expression may enhance actin polymerization in the TM. Furthermore, the permeability of the TM cell layers was inhibited by RHOB overexpression (*p* < 0.01, unpaired t-test, Fig. 5O) and dobutamine treatment (*p* < 0.05 vs. DMSO, Šidák multiple comparison test, Fig. 5P), whereas RHOB KO canceled this inhibition (Fig. 5P). Taken together, these suggest that NE mainly inhibits TM phagocytosis to decrease AH permeability through RHOB-ROCK activation via the β1AR-cAMP-CREB signaling pathway.

### Nocturnal activation of β1AR-ROCK increases IOP

Since NE in the AH has a circadian rhythm with a nocturnal increase in rodents^51^, which seems to also be true in humans, a nocturnal increase in NE in the TM may cause an IOP increase by increasing RHOB. To test this hypothesis, we administered Y27632 and RKI1447 (ROCK inhibitors) and rhosin (RHO inhibitor) to mouse eyes at ZT10 and measured the nighttime IOP (ZT15) (Fig. 6A–C). Y27632 and RKI1447 (replot data^17^) administration at ZT10 arrested the increase in nocturnal IOP; in particular, ROCK inhibitors significantly suppressed IOP (Fig. 6B). At the individual level, these two and rhosin administration suppressed the nocturnal IOP increase (Fig. 6C). ROCK inhibitors, which block the nocturnal IOP increase in mice^17^, are widely used for glaucoma therapy^21–23^. These results indicate the role of RHO and ROCK in the inhibition of TM phagocytosis and the increase in cell adhesion during the nocturnal increase in IOP. To examine their involvement in daytime IOP reduction, we administered Y27632, and RKI1447, rhosin, and LPA to mice at ZT4 and measured IOP at ZT9 (Extended Data Fig. 10A,B). LPA treatment significantly increased IOP, but not the other inhibitors (*p* < 0.01, unpaired t-test, Extended Data Fig. 10A). Individual data also showed a tendency for LPA to increase the diurnal IOP (Extended Data Fig. 10B), indicating that daytime activation of RHO may increase the IOP. Nocturnal IOP suppression by ROCK and RHO inhibitors was similar to a previous report demonstrating that RhoA blocking by adeno-associated virus prevents nocturnal IOP elevation in rats^52^. Taken together, these findings suggest that AH outflow regulation by the cytoskeleton is night-limited or time-independent, whereas TM phagocytosis may mediate the daytime increase in AH outflow.

**Fig. 6:**
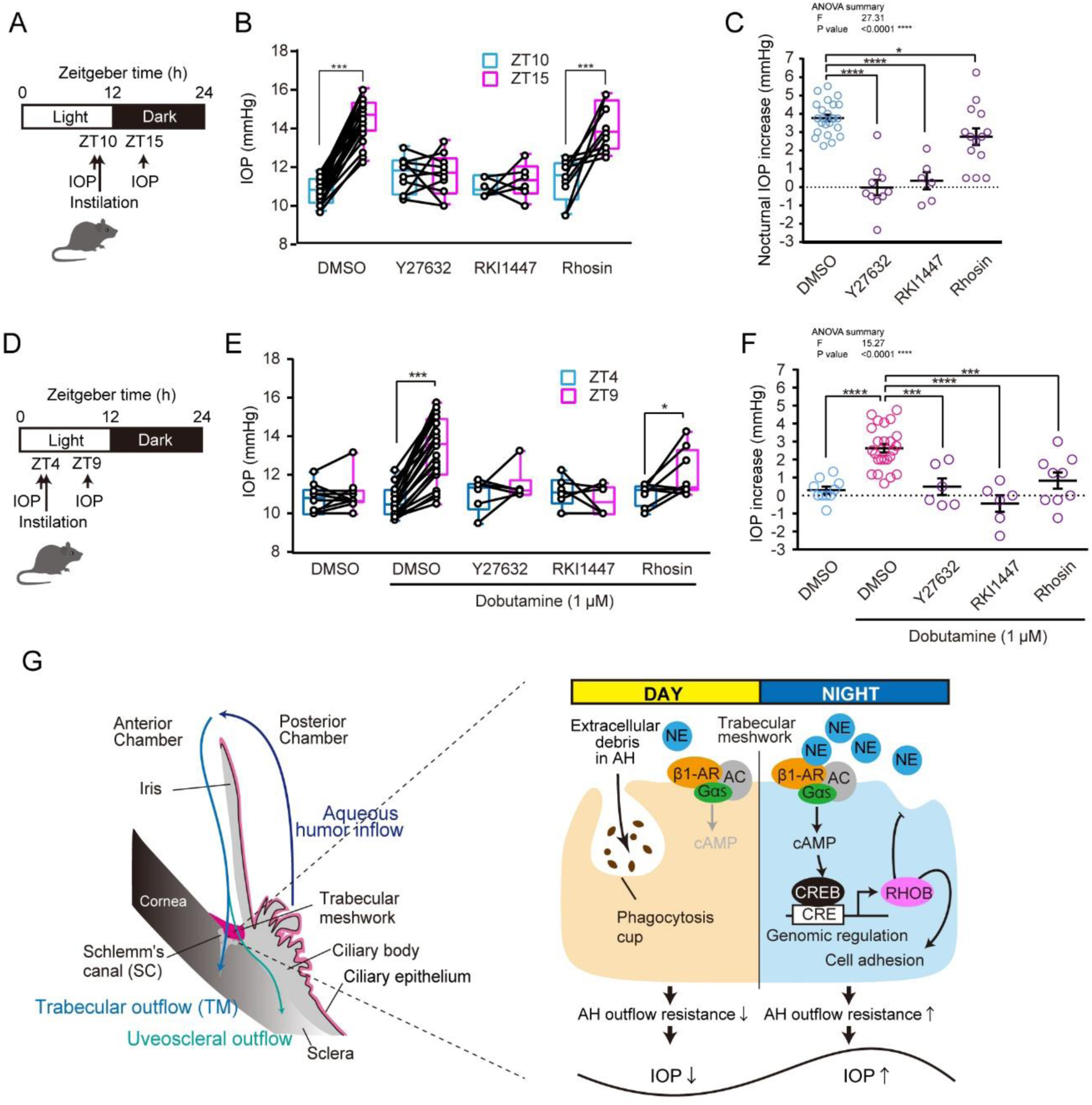
Norepinephrine (NE) induced nocturnal increase in intraocular pressure (IOP) via the RHO-ROCK1 pathway. **A–C** Effect of RHO-ROCK pathway-related inhibitors on nocturnal IOP increase in mice. Mice were administered Y27632 (100 µM/0.1% DMSO PBS), and rhosin (100 µM) at ZT10, and IOP was measured at ZT15. **B** Y27632 and rhosin treatment significantly suppressed the nocturnal IOP increase (paired t-test, \*\*\**p* < 0.001). Data are presented as box-and-whisker plots of individual data (n = 6–16). **C** Inhibitors prevented IOP increase (\**p* < 0.05, \*\*\*\**p* < 0.0001 vs. DMSO, one-way ANOVA [*p* < 0.001], Dunnett’s multiple comparison test). Data are presented as scatter plots with mean ± SEM (n = 6–16). **D,E** Validation of the inhibitory effects of drugs on IOP using dobutamine (100 µM) treatment at ZT4. After 5 h, IOP was measured. Dobutamine significantly increased IOP, whereas pretreatment with Y27632 and rhosin prevented this effect (paired t-test, \**p* < 0.05, \*\*\**p* < 0.001). **F** The same was true for individual data (\*\*\**p* < 0.001, \*\*\*\**p* < 0.0001 vs. Dobutamine+DMSO, one-way ANOVA [*p* < 0.0001], Dunnett’s multiple comparison test). Data are presented as (**E**) box-and-whisker plots with individual data and (**F**) as scatter plots with mean ± SEM (n = 6–16). **G** Regulatory model of time-dependent systems by which nocturnal NE suppresses phagocytosis to induce AH outflow resistance through β1AR-cAMP-CREB-RHOB-ROCK activation, leading to an IOP increase at night.

The role of the β1AR-RHOB-ROCK pathway in diurnal IOP changes remains unclear. Since daytime dobutamine administration elevates IOP in mice^17^, we administered Y27632, RKI1447, and rhosin with dobutamine to mice at ZT4 and measured IOP at ZT9 (Fig. 6D–F). Administration of ROCK inhibitors (replot of RKI1447^17^) significantly suppressed the nocturnal IOP increase (Fig. 6E) at the individual level (*p* < 0.01 vs. DMSO, Dunnett’s multiple comparison test; Fig. 6F), suggesting a role for the β1AR-RHOB pathway in nocturnal IOP increase. Since NE from the SCG peaks during nighttime in rodents^53^, the circadian rhythm of SCG-NE may regulate diurnal changes in TM phagocytosis (Fig. 6G). Taken together, these findings show that nocturnal NE suppresses phagocytosis-mediated AH outflow through β1AR-RHOB activation, leading to an increase in nocturnal IOP (Fig. 6G).

## Discussion

Previous studies have demonstrated that sympathetic NE transmits circadian signals to the eye to generate IOP rhythm^13^, while β1AR suppresses TM phagocytosis by silencing PIP3-AKT or ERK signaling through the cAMP-EPAC-SHIP1^16^. However, the role of NE in regulating TM phagocytosis via gene expression remains unknown^14,15^. In this study, we found that NE contributes to the elevation of nocturnal IOP through genomic regulation involving β1AR-cAMP-CREB-RHOB-ROCK activation (Fig. 1–3). A recent microarray analysis identified β1AR blocker betaxolol as the top compound that opposed POAG signatures^54^. Activation of cAMP signaling pathway and CREB was observed in the TM of POAG donors^55^. Moreover, β1AR stimulation was found to enhance cAMP-CREB signaling in TM cells^17^, supporting the results of the present study. Furthermore, although NE suppressed TM phagocytosis, it promoted actin polymerization, increased cellular adhesion, and accelerated TM cell aging (Fig. 4). NE-induced RHOB elevation in TM cells primarily contributed to the inhibition of macrophage phagocytosis (Fig. 5). While RHOA typically promotes phagocytosis in TM cells through F-actin formation^56^, RHOB has been implicated in hypoxia-induced macrophage phagocytosis^57^; however, its role in TM macrophage phagocytosis remains unclear. The present study showed that RHOB suppresses phagocytosis in TM cells (Fig. 5). Ocular RHO or ROCK inhibition in mice prevented both nocturnal and NE-induced increases in IOP (Fig. 6). This aligns with previous reports demonstrating the IOP-lowering effects of ROCK inhibitors^21–23^. Since the detailed mechanisms remain to be clarified, a further understanding of this pathway is essential to elucidate the complex cellular mechanisms by which AR activation regulates aqueous humor dynamics, potentially contributing to the development of chronotherapy strategies.

Because of the long-term effect (more than 4h) on RHOB activity after NE instillation (Fig. 2), we expected that NE increases RHOB activity by protein level in the TM. However, RHO GTPase activity acts on many important molecular switch in several cells and is strictly regulated by RHO Guanine-nucleotide exchange factor, GDP-dissociation inhibitor (GDI), and GTPase activating protein. However, we could not detect significant changes in these genes in upregulated 18 genes (Fig. 1), indicating the absence of regulatory mechanisms of RHOB activationa by a specific gene. Interestingly, GSEA in molecular function revealed significant GTPase related terms in the TM cells (Fig. 4). It is conceivable that there may be an integrated regulatory mechanism for GTPase activity rather than a regulatory mechanism for RHO activity by specific gene expression.

We found that RHOB in TM macrophages inhibited phagocytic activity and slightly promoted cell adhesion (Fig. 5). Several studies have demonstrated that RHOB plays a role in the inflammatory response, particularly in macrophages and endothelial cells. RHOB is involved in mannose receptor-mediated phagocytosis in human alveolar macrophages^64^. RHOB influences macrophage adhesion and migration by reducing the surface expression of β2 and β3 integrins. However, this is not essential for podosome formation, as shown in experiments using primary macrophages from *RHOB*-deficient mice^65^. *RHOB*-deficient macrophages exhibit enhanced motility^65^, which correlates with increased recruitment during peritoneal inflammation *in vivo*^66^. Despite these roles, RHOB does not affect actin stress fiber formation, cell growth, or cellular senescence in the TM (Fig. 5), whereas RHOA/ROCK activation induces actin filament formation in TM cells^21^. The RHOB protein sequence is highly similar to RHOA and RHOC; however, it exhibits distinct biochemical and biological properties. Although all three proteins undergo C-terminal prenylation, RHOA and RHOC are exclusively geranylgeranylated, whereas RHOB is either geranylgeranylated or farnesylated. In addition, only RHOB undergoes palmitoylation^67,68^. Unlike RHOA and RHOC, which predominantly localize in the cytoplasm through interaction with RHO GDI, RHOB does not bind to RHO GDI, but instead localizes to the plasma membrane and/or endosomes^44^. RHOB plays a crucial role in regulating cell cycle and apoptosis, and its expression declines during aging and tumorigenesis but is upregulated in response to DNA damage. This suggests its potential involvement in regulating aging-related cellular senescence, oncogene-induced senescence, and stress-induced senescence (e.g., caused by irradiation), warranting further investigation.

This study is the first to demonstrate that NE induces RHOB expression via the cAMP-CREB-CRE pathway in mammals. Although RHOB expression was previously shown to be regulated by ultraviolet irradiation, ERK, AKT, and Rho, its regulation through ARs remained unknown^33,34,62^. Since NE suppresses AKT activation in human TM cells^17^ and the PI3K/AKT pathway is known to downregulate RHOB expression^63^, it is possible that RHOB is also regulated by AKT-mediated NE signaling. Conversely, β1AR agonists such as L-NE and dobutamine stimulate cAMP-CREB activation in iHTMCs^17^. No CRE sites have been identified in the promoters of RHOA or RHOC^33,34^. Furthermore, according to the Signaling Pathways Project database (http://www.signalingpathways.org/ominer/query.jsf), CRE was not found to induce RHOA expression. This suggests that RHOB-specific expression may be mediated by CRE.

In mice, the IOP-lowering effect of ROCK inhibitor eye drops was greater than that of the RHO inhibitor treatment (Fig. 6). This suggests that pathways other than RHOB may contribute to the regulation of AH outflow by NE or GCs. Although the exact concentration of the inhibitors was not verified, ROCK inhibition is likely more effective for the rearrangement of the actin cytoskeleton and softening of TM. Moreover, cAMP/PKA activation and subsequent RHOA inactivation result in the loss of actin stress fibers, focal adhesion disassembly, and disruption of the extracellular matrix network in the TM^29^, implying that the ROCK pathway plays a role in IOP suppression independent of NE-induced RHOB signaling. The effects of actin polymerization inhibitors on IOP reduction in mice appear to be restricted to nighttime^16^. During daytime, suppression of AH outflow due to reduced TM phagocytosis seems to outweigh the effects of actin cytoskeletal rearrangement^16^. The cellular circadian clock regulates various cytoskeletal regulators in fibroblasts^69^. In mice, approximately 30–42% of AH passes through the uveoscleral pathway^70^. To fully elucidate the mechanisms governing the rhythmic dynamics of AH outflow, these factors must be investigated further.

The circadian regulation of NE/GC by key factors such as cytoskeletal dynamics, cell adhesion, and IOP-independent uveoscleral outflow to the ciliary muscle remains poorly understood. In the present study, NE inhibited cell proliferation and growth in a dose-dependent manner, whereas low doses induced cellular senescence (Fig. 5). In addition, NE promoted actin polymerization and cell adhesion (Fig. 5). In contrast, Dex stimulated cell growth, actin polymerization, and cellular senescence in a dose-dependent manner (Extended Data Fig. 7). Notably, both NE and Dex induced actin polymerization and enhanced cell adhesion, which contributed to increased TM stiffness and increased AH outflow resistance. Both β-adrenergic signaling and GCs mediate SCN timing signals in osteoblasts^58^. Previous studies have reported interactions between the sympathetic nervous system and GCs, with GCs transcriptionally modulating β2-AR expression by regulating GC-responsive elements on its promoter^59^. Interestingly, GCs can rapidly activate cAMP production via Gαs to initiate non-genomic signaling, which contributes to approximately one-third of their canonical genomic effects^60^. Supporting these findings, betaxolol has been shown to prevent steroid-induced IOP elevation^61^. Therefore, in the TM, Gs-bound glucocorticoid receptors may enhance β1AR-Gs signaling, modulate TM stiffness, and regulate the AH drainage rhythm.

Our study had several inherent limitations. First, the use of the pan-RHO inhibitor rhosin in this study may not have fully explained the specific contribution of RHOB. Aithough *RHOA* and *RHOC* were not upregulated by NE, these RHOs may not have NE-mediated TM phagocytosis suppression. TM-specific KO using viral delivery *in vivo* may provide further insights into the role of RHOB in IOP regulation. Second, although we proposed a model for IOP induction driven by nocturnal NE, the circadian variation in NE release from the SCG in humans remains unclear. While GC secretion peaks at the light offset^71^ and SCG-NE shows a nocturnal peak in rodents^53^, GC rhythms are anti-phasic to NE rhythms in diurnal animals^53,72^. In humans and other mammals, nocturnal NE release from the SCG typically stimulates melatonin synthesis^73^. β-blockers in humans have been shown to inhibit nocturnal melatonin levels and suppress IOP elevation from late night to early morning^74^. Similarly, in nocturnal rabbits, β-blockers suppress IOP increase only at night^75^. These findings suggest the role of nocturnal NE release from the SCG in regulating IOP rhythms. Third, our current model cannot fully explain the differences in IOP rhythms between diurnal and nocturnal animals. In nocturnal animals, IOP peaks early at night^7^, whereas in healthy humans, it remains elevated during the night and peaks between late night and early morning^6,9^. In phasic SCG-NE and anti-phasic GC, the action of both factors on the IOP increase may explain these differences.

In summary, these findings indicate a potential circadian function for the RHOB-mediated NE regulation of phagocytosis and AH outflow resistance in the TM, thereby influencing the IOP rhythm. Although the TM is responsible for the majority of AH outflow and serves as the primary site of outflow resistance^70,76^, no clinically approved drug had a direct impact on TM until the advent of ROCK inhibitors. Furthermore, there are no established IOP rhythm regulators. Although innovative treatments with new mechanisms, such as chronotherapy, are critically needed for glaucoma management, this study proposes the potential for developing therapeutic agents that target RHOB as an IOP rhythm regulator. Since genomic signaling might also contribute to IOP rhythm regulation^60^, further understanding of the time-dependent effects of NE and GC on AH dynamics is essential to fully elucidate the regulatory mechanisms of IOP rhythm.

## Methods

### Animals

Five-week-old male C57BL/6JJmsSlc mice (N =130; Japan SLC Inc., Shizuoka, Japan) were purchased and housed in plastic cages (170 W × 240 D × 125 H mm; Clea, Tokyo, Japan) under a 12-h light (200 lx of fluorescent light)/dark cycle (12L12D, 0800 light ON, 2000 light OFF), and maintained at a constant temperature (23 ± 1 °C). Food (CE-2; CLEA) and water were provided *ad libitum*^13^. All animal experiments were conducted in accordance with the Guidelines for Animal Experiments of Aichi Medical University and the Faculty of Agriculture at Kyushu University. All experiments were approved by the Animal Care and Use Committee of Aichi Medical University and Kyushu University.

### IOP measurement

IOP measurements were performed using a tonometer (Icare TonoLab, TV02; Icare Finland Oy, Espmoo, Finland) as previously reported^8,13^. All mice were maintained under 12L:12D conditions for more than two weeks before IOP measurements. The unanesthetized mice were gently held using a sponge. IOPs were measured during the light phase under light (200 lx) conditions and during the dark phase under dim red light conditions. For the analysis of RHO-related drugs and dobutamine-mediated IOP induction, IOP was measured before drug administration at zeitgeber time (ZT) 4 and measured at ZT9. ZT0 (0800) was defined as the time at which the light was ON. To analyze the nocturnal increase in IOP, IOP was measured at ZT10 and ZT15. We calculated back from ZT15 (the peak of IOP) for 5 h and set it to ZT10 when the IOP was low.

### Drug administration

Drugs were administered as described in our previous report^13^. Unanesthetized 8-week-old male mice were used in this study. For microarray analysis, 7 weeks old male mice (N = 6) were adrenalectomized and superior cervical ganglionectomized, as in our previous report^13^. Two weeks after, mice were treated with drugs at ZT2, when IOP is low, with a single drop (30 μL) of L-NE (1 mM/0.1% DMSO PBS) or dobutamine (1 mM/0.1% DMSO PBS) using a micropipette into bilateral eyes. During instillation, the mice were gently restrained with their necks held back. After 4 h, total RNA of vehicle- or NE-treated eyes was extracted using the DNA/RNA/Protein Extraction Kit (#DRP100, Presto) at ZT6 and pooled for microarray analysis (N = 1). Six hours after dobutamine instillation, proteins were extracted from the eyes using cell lysis buffer (#9803, Cell Signaling Technology, Tokyo, Japan) containing a protease inhibitor cocktail (#P8340, Sigma Aldrich) and phosphatase inhibitor cocktail 1 (#P2850, Sigma Aldrich) according to the manufacturer’s instructions.

To analyze the inhibitory efficacy of RHO-related drugs on dobutamine-induced IOP increase, mice were pretreated at ZT4 with a single drop (30 μL) of ROCK1/2 inhibitor RKI1447 (1 mM/0.1% DMSO PBS; Cayma, 16278) or Y27632 (1 mM/0.1% DMSO PBS; FUJIFILM WAKO, 030-24021), or pan RHO inhibitor rhosin (1 mM; FUJIFILM WAKO, 5003101) using a micropipette into bilateral eyes. Then, after 10 min, a single drop of dobutamine (100 µM/0.1% DMSO PBS) was added. During drug administration, the mice were gently restrained with their necks held back. After 5 h, the IOP was measured at ZT9. For nocturnal IOP increase analysis, mice were treated with a single drop of the above drugs at ZT10 using a micropipette in both eyes, and IOP was measured at ZT15.

### iHTMC culture

The immortalized human trabecular meshwork cell line (iHTMC) TM-SV40 derived from primary human SC and TM regions was purchased from Applied Biological Materials Inc. (T-371-C, ABM Inc., Richmond, BC, Canada) and cultured in TM cell medium (TMCM; #6591, Sciencell, Carlsbad, CA, USA) supplemented with 1% fetal bovine serum (#0010, ScienCell), 1% growth supplement (TMCGS, #6592, Sciencell) and 1% penicillin/streptomycin (#0503, Sciencell) in a type I collagen-coated 100-mm dish (#3020-100, IWAKI, Japan). The experiments were performed on type I collagen-coated plates. Upon reaching confluence, iHTMCs were split 1:3 using 0.05% trypsin/PBS. Cell viability was determined using trypan blue (0.4%) exclusion assay.

### Transcriptome microarrays analysis

Genome-wide transcriptome analysis was performed using Affymetrix GeneChip™ Whole Transcript (WT) PLUS Reagent Kit and GeneChip™ Human Clariom™ D Arrays according to the manufacturer’s instructions (ThermoFisher). Total RNA (250 ng) was used as the starting material for target preparation. Microarrays were subsequently washed, stained by GeneChip™ WT PLUS Reagent Kit, and scanned using GeneChip™ Scanner 3000 7G and the Affymetrix GeneChip™ Command Console Software with .cel files as data output by FilGen. The Microarray Data Analysis Tool (Filgen, Inc., Japan) facilitated the interpretation of gene expression data generated by the Clariom D microarrays.

### RNA-seq analysis

High–quality total RNA was extracted using a DNA/RNA/Protein Extraction Kit (#DRP100; Presto) and verified using an Agilent 2100 Bioanalyzer. RNA-seq was performed by GENEWIZ using DNBSEQ with a read configuration of 150 bp for paired-end reads, and 40 million reads were generated per sample. The obtained FASTQ files were analyzed by RaNA-seq with the DESeq2 package using the non-paired Wald parametric method to obtain counts and transcripts per million (TPM) data. RaNA-seq is an open bioinformatics web tool for quick analysis and visualization of RNA-seq data (available at http://ranaseq.eu). It performs a full analysis in minutes by quantifying FASTQ files, calculating quality control metrics, running differential expression analyses, and explaining the results with functional analyses^77^. The adjusted p value cutoff was set at 0.05.

### Enrichment analysis

To identify the important terms related to 18 genes upregulated by NE stimuli, we performed Gene Ontology (GO) enrichment analysis for molecular function and biological processes linked by adjusted p values using the web tool Enrichr (https://maayanlab.cloud/Enrichr/) and created graphs using the web tool SRplots (https://www.bioinformatics.com.cn/en).Among the reactome, kinase enrichment analysis for kinase perturbations from GEO down and GEO up, and transcription factor (TF) enrichment analysis for ENCODE and ChEA Consensus TFs from ChIP-X, the important terms were determined depending on the combined enrichment score. Gene set enrichment analysis (GSEA) for molecular function, biological process, and cellular components of NE-treated iHTMC was conducted using RaNA-seq (https://ranaseq.eu/index.php) and graphs were created using the web tool SRplots. Terms were launched based on normalized enrichment scores.

To find RHOB-related transcriptome data, we used publicly available data from The Signaling Pathways Project (http://www.signalingpathways.org/ominer/query.jsf) in addition to our NE-treated human TM cells data (226 upregulated genes by 6 h NE exposure to iHTMC [PRJDB20330; PADJ < 0.05]); 75 upregulated genes in human primary osteoblasts after 2 h exposure of prostaglandin E2 (PGE2)-activating GPCR (EP2 or EP4 receptor) (GSE21727; FDR < 0.05, Log2FC > 2), 130 upregulated genes in human K562 leukemia cells by CREB1 knockdown (GSE12056; PADJ < 0.05 Log2FC > 2), 571 upregulated genes in rat liver 3 h after injection with Gi-coupled α2AR blocker chlorpromazine (25313160; PDR < 0.05 Log2FC > 2), and 3,386 upregulated genes in mouse immortalized preadipocyte after 4 h of dibutyryl-cAMP exposure (GSE5041; FDR < 0.05 Log2FC > 1). SR plots was used to create a graph of the upset plot for the five datasets.

### Comparative analysis of RNA-seq and microarray data

To compare the overlap of differentially expressed genes in the two statistical comparisons, the Venn diagram web tool (https://bioinformatics.psb.ugent.be/webtools/Venn/) was used to identify 18 commonly upregulated genes. SRplot (https://www.bioinformatics.com.cn/en) was used to create scatter plots of microarray and RNA-seq data, cluster heatmaps of RNA-seq in iHTMC, and correlation matrices of Pearson coefficients of GO analysis of the 18 genes^78^.

### RHOB activity assay

To measure iHTMC RHOB activity in a collagen-coated 6-well plate (#636-35533, IWAKI), the RhoA G-LISA GTPase Activation Assay Kit (colorimetric format) (BK124-S, Cytoskeleton, Inc.) was used according to the manufacturer’s instructions, and modified for RHOB activity measurement by using RHOB antibody as previous report^79^. Briefly, 6 h after dobutamine administration, the culture plate was retrieved from the incubator, immediately placed on ice, the medium was aspirated, and the cells were washed twice with 1 ml ice-cold PBS. iHTMCs were lysed for 15 min in ice-cold cell lysis buffer, 50 μl per well. After centrifuging cell lysates at 10,000 × *g*, 4 °C for 10 min, total supernatant protein concentrations were quantified using a TaKaRa BCA Protein Assay Kit (T9300A, Takara). The supernatant was stored at −80 °C. We kept a number of strips of Rho plate on ice and added 100 μL ice-cold water in each well to dissolve the powder coat, covering the bottom of the well. Then, we added 5 equalized cell lysates to wells and placed the plate on an orbital microplate shaker (250 rpm) at 4 °C for 30 min. Active, GTP-bound Rho in cell lysates will bind to the wells while inactive GDP-bound Rho is removed during washing steps. After washing twice with a wash buffer, antigen-presenting buffer was added to each well and incubated at room temperature for 2 min. The antigen-presenting buffer was flicked out vigorously, and the plate was washed three times with wash buffer. We added 50 µL of diluted anti-RhoB primary antibody (1:100; 14326-1-AP, ProteinTech) to each well and left the plate on an orbital microplate shaker (250 rpm) at room temperature for 45 min. After washing, secondary horseradish peroxidase (HRP)-labeled anti-rabbit antibody (1:62.5, 7074, Cell Signaling Technology) diluted in Antibody Dilution Buffer was added to each well, and the plate was incubated on a microplate shaker (250 rpm) at room temperature for 45 min. After incubation with mixed HRP detection reagent at 37 °C for 10 min, we calculated the signal by measuring absorbance at 490 nm using a Nivo microplate reader.

### Western blot analysis

Western blotting was performed as previously described^80^. Protein extraction was performed using a cell lysis buffer (9803, Cell Signaling Technology, Tokyo, Japan) containing a protease inhibitor cocktail (P8340, Sigma Aldrich) and phosphatase inhibitor cocktail 1 (P2850, Sigma Aldrich) according to the manufacturer’s instructions. The total protein transferred to a polyvinylidene fluoride membrane and was detected using EzStainAQua MEM (WSE-7160, ATTO) and used for normalization. After destaining, membranes were incubated with rabbit polyclonal antibody against RHOB (1:1,000; 14326-1-AP, ProteinTech). Membranes were washed and incubated with HRP-conjugated goat polyclonal antibodies against mouse and rabbit IgG (1:10,000; 7074, Cell Signaling Technology). Chemiluminescent images were obtained using FUSION-SOLO.7S.EDGE (Vilber Bio Imaging).

### *RHOB* promotor assay

Since no previous studies have reported the presence of CRE on *RHOB* promoter in any animals, which bind CREB or CREB family members^36^, annotation analysis of human *RHOB* promotor (−1,500 bp to +1 bp) using UCSC genome browser revealed that not only highly conserved TATA box (TATATTAA; −56 bp to −49 bp) and CAAT element (CCAAT; −115 bp to −111 bp) but also one previously unknown CRE′ site (TGAGCTCA; −89 bp to −82 bp) near the transcription start site in mammals. To investigate directly the role of this CRE in NE-mediated increase in *RHOB* promoter activity (−118 bp to +44 bp), we designed and purchased predesigned a short human *RHOB* promotor sequence (−119-+44; wild-type (WT) CRE′ [TGAGCTCA]) plasmid (pRP[Pro]- hRluc/Puro-{human *RHOB* promoter −119-+44}>TurboGFP; Vector builder) and built a series of plasmid containing reporter constructs with WT CRE′, normal CRE (TGACGTCA) by reversing the middle GC, and random CRE (CCAAGGTT). We seeded iHTMCs in 2 mL of serum (+) TMCM in a collagen-coated 6-well plate (#636-35533, IWAKI) 1 d before transfection to achieve 50–70% confluency at the time of transfection. We diluted 1 µg of plasmid DNAs in 50 µL of Xfect Reaction Buffer (Takara Bio, # 631318), and then added 0.3 µL of Xfect Polymer to the diluted DNA. After incubation for 10 min at room temperature to form the transfection complex, the prepared transfection complex was gently added dropwise to each well. After 4 h incubation at 37 °C in a humidified incubator with 5% CO₂, we replaced the transfection medium with fresh growth medium and incubated to recover and express the neomycin resistance gene for 48 h. After 48 h post-transfection, we replaced the medium with fresh growth medium containing 1 µg/mL of puromycin (FUJIFILM WAKO) for continuous selection for 3 d, changing the medium every 2 d, until untransfected cells are eliminated. After checking bioluminesence by Renilla luciferase (Rluc) with the substrate Coelenterazine 400a (#300-1, NanoLight technology) using a microplate reader Nivo (Perkin Elmer), we monitored TurboGFP expression after L-NE (1 μM) stimuli in iHTMC using a microplate reader Nivo (Ex 480 nm, Em 530 nm; Perkin Elmer).

### RHOB knock-out

To knock out *RHOB* in iHTMC, we designed a specific sgRNA targeting the human *RHOB* gene-cloned CRISPR/Cas9 plasmid (pRP[2CRISPR]-EGFP/Neo-hCas9-U6>hRHOB[gRNA#109]-U6>hRHOB[gRNA#139]; Vector builder). We seeded iHTMCs in 2 mL of serum (+) TMCM in a collagen-coated 6-well plate 1 d before transfection to achieve 50–70% confluency at the time of transfection. We diluted 1 µg of plasmid DNAs in 100 µL of Xfect Reaction Buffer (Takara Bio, # 631318), and then add 0.3 µL of Xfect Polymer to the diluted DNA. After 10 min incubation at room temperature to form the transfection complex, the prepared transfection complex dropwise was gently added to each well containing 1 mL of serum (+) TMCM. After 4 h incubation at 37 °C in a humidified incubator with 5% CO₂, we replaced the transfection medium with fresh growth medium and incubated to recover and express Neomycin resistance gene for 48 h. After 48 h post-transfection, we replaced the medium with fresh growth medium containing 700 µg/mL of G418 (158782, Fujifilm Wako) for continuous selection for 11 d, changing the medium every 2 d, until untransfected cells are eliminated. After checking the EGFP expression using a fluorescence microscope, we detected the absence of RHOB protein in iHTMC using western blot analysis.

### Liquid permeability assay

To verify the liquid permeability of iHTMC multilayer culture in culture inserts (for 24 well with 0.4 μm PET, Falcon; #353095), we employed a multilayer culture to mimic the TM structure which returned the color of the culture medium to the inside and outside of the culture insert to quantify the coloration of the medium after 2 h using an absorbance system. A high density of iHTMC was cultured on a culture insert PET membrane with serum-(+)-TMCM for 2 d. iHTMC multilayer formation was observed non-invasively via trans-epithelial electrical resistance measurements using a Millicell-ERS2 (Electrical Resistance System; #MERS00002, Millipore). Barrier formation was observed with an increase in the resistance. After the medium was replaced with serum-free TMCM, L-NE was added for several exposure times (0, 1, 2, 4, 6, 8, and 24 h before measurement). After removing the medium, phenol-red-positive Dulbecco’s Modified Eagle Medium (DMEM; 600 μL, #D6429, Sigma Aldrich) was added to culture inserts while the phenol red-free DMEM was added outside of inserts (600 μL). After 2 h, the outside medium (100 μL) was transferred to a new 96-well plate (Thermo Fisher) to measure absorbance (560 nm) using a Nivo microplate reader. To investigate the effect of RHOB KO, KO iHTMC were cultured and exposed to dobutamine for 6 h. as described above.

### Phagocytosis assay

For the phagocytosis assay, WT and RHOB KO iHTMCs were plated in collagen I-coated 96-well microplates (4860-010; IWAKI) at a density of 5.0 × 10^3^ cells/well in TMCM supplemented with 1% penicillin/streptomycin and growth factors (6591; Sciencell). To measure phagocytosis, pHrodo Green Zymosan Bioparticles (P35365; Thermo Fisher Scientific) were suspended in TMCM and vortexed to disperse. After 90% confluence, the medium was removed by aspiration, and 100 μL of serum-free TMCM was immediately added. After 24 h, the medium was replaced with serum-free TMCM containing pHrodo Zymosan (2.5 µg/well) in the presence of several kinds of drugs, and the plate was placed in a 5% CO_2_ incubator at 37 °C. No pHrodo zymosan was used as fluorescent control using background fluorescent intensity because of autofluorescence in TMCM, and vehicle control included 0.1% DMSO. To confirm the effects of RHOB KO, the green fluorescence intensity in each well was measured using a Nivo microplate reader (Ex 495 nm/Em 530 nm, PerkinElmer, #HH3500) 6 h after dobutamine (Sigma-Aldrich, #D0676) administration and 4 d after Dex (Dexamethasone, #11107-51, Nakarai Tesk, Kyoto, Japan) administration. Subsequently, at the time of medium replacement, Dobutamine and Dex were added to the serum-(-) TMCM at final concentrations of 0, 0.001, 0.01, 0.1, 1, and 10 µM. The phagocytosis assay was independently repeated thrice using three or four biological replicates.

To analyze the inhibitory efficacy of RHO-related drugs on dobutamine-induced phagocytosis suppression, iHTMCs were treated using Y27632 (0.1, 1, and 10 µM), or rhosin (0.1, 1, and 10 µM) with dobutamine (1 µM). To confirm the effect of RHO activator lysophosphatidic acid (LPA; 1 mM) on TM phagocytosis, LPA (0.01, 0.1, 1, and 10 µM) were added to iHTMC.

### Actin polymerization assay

After 90% confluence of WT and RHOB KO iHTMCs, the medium was removed by aspiration, and 100 μL of serum-free TMCM was immediately added. After 24 h, the medium was replaced with serum-free TMCM containing 100 nM SaraFluor 497 actin probe (AR5601-N5, Goryo Chemical, Inc., Japan) to stain actin filaments under several concentrations (0.01, 0.1, 1, and 10 µM) of L-NE. The plate was placed in the IncuCyte ZOOM instrument (Essen Bioscience, Ann Arbor, MI, USA), and incubated in a 5% CO_2_ incubator at 37 °C. Each well was imaged at 3 points, every 0.5 h or 1 h for more than 62 h using the channels of green fluorescence and phase and the 10× objective (Nikon). No probe was used for fluorescent control using background fluorescence intensity because of autofluorescence in TMCM, and vehicle control included 0.1% DMSO. The green fluorescence intensity at each time point in each well was measured using IncuCyte ZOOM 2015A software (Essen Bioscience).

To confirm the effects of RHOB KO, fluorescence (Ex 480 nm/Em 530 nm) was measured using a Nivo microplate reader (PerkinElmer, #HH3500) 6 h after dobutamine administration and 4 d after Dex administration. Subsequently, at the time of medium replacement, Dobutamine and Dex were added to the serum-(-) TMCM at final concentrations of 0, 0.001, 0.01, 0.1, 1, and 10 µM.

### Cell growth

WT and RHOB KO iHTMCs were plated in collagen I-coated 96-well microplates (#4860-010; IWAKI) at a density of 2.5 × 10^3^ cells/well in TMCM supplemented with 1% penicillin/streptomycin and growth factors (#6591; Sciencell). After 48 h, 10 μL of cell counting kit-8 reagent (CCK-8, #CK04, Dojindo Laboratories) was added to cells seeded in 96 wells, and incubated for 1 h in a 37 °C CO_2_ incubator. To calculate cell numbers, iHTMC were preincubated in collagen I coated 96-well microplates (4860-010; IWAKI) at densities of 16 × 10^3^, 8.0 × 10^3^, 4.0 × 10^3^, 2.0 × 10^3^, 1.0 × 10^3^, and 0.5 × 10^3^ cells/well with serum free TMCM (1% penicillin/streptomycin; #6591; Sciencell), and were counted as standard. Subsequently, the absorbance (450 nm) was measured using a Nivo microplate reader (PerkinElmer, #HH3500).

Subsequently, at the time of medium replacement, Dobutamine and Dex were added to the serum-(+) TMCM at final concentrations of 0, 0.001, 0.01, 0.1, 1, and 10 µM. The medium was replaced with fresh medium. The effect of these compounds on the cell proliferation rate was assessed using the Cell Counting Kit-8.

### Cell adhesion assays

WT and RHOB-KO iHTMCs were plated in collagen I-coated 96-well microplates (#4860-010; IWAKI) at a density of 5 × 10^3^ cells/well in serum (+) TMCM (#6591; ScienCell). After reaching 100% confluency, the medium was changed to serum-free TMCM for 24 h. Then, dobutamine and Dex were added to the serum (+) TMCM at final concentrations of 0, 0.001, 0.01, 0.1, 1, and 10 µM. After 8 h, iHTMCs were washed once with PBS, detached with 30 uL of 0.25% trypsin treatment for 5 mins and diluted with 170 µL of serum (+) TMCM. The entire volume was seeded onto new collagen-coated 96-well plates (#4860-010; IWAKI), placed in a CO_2_ incubator for 2 h, washed twice with PBS, and adherent cells were quantified as described above. The adhesion rate was calculated as the ratio of the absorbance of the treated samples to that of the control.

After 72 h of fluorescent measurement, 10 μL of cell counting kit-8 reagent (CK04, Dojindo) was added to cells seeded in quadruplicate in 96 wells and incubated for 4 h in a 37 °C CO_2_ incubator. To calculate cell numbers, iHTMC were preincubated in collagen I coated 96-well microplates (4860-010; IWAKI) at densities of 16 × 10^3^, 8.0 × 10^3^, 4.0 × 10^3^, 2.0 × 10^3^, 1.0 × 10^3^, and 0.5 × 10^3^ cells/well with serum free TMCM (1% penicillin/streptomycin; #6591; Sciencell), and were counted as standard. Subsequently, the absorbance (450 nm) was measured using a Nivo spectrophotometer (PerkinElmer).

For temporal analysis of iHTMC adhesion according to NE exposure time, we fitted a logistic growth curve using GraphPad Prism 8.

Y=YM*Y0/((YM-Y0)*exp(-k*x) +Y0)

where Y0 denotes the starting population (same units as Y), YM represents the maximum population (in the same units as Y), And k is the rate constant (the inverse of X). 1/k is the x-coordinate of the first inflection point.

### RHOB overexpression

We seeded iHTMCs in 2 mL of serum (+) TMCM in a collagen-coated 6-well plate 1 d before transfection to achieve 50–70% confluency. To express RHO in iHTMC, we designed pcDNA3.1+ plasmids containing nano-luciferase (Nluc)-conjugated to the N-terminus of wild-type (WT) RHOB and RHOC cloned by standard PCR protocol^81^ from the cDNA of human melanoma cell line, Mel270, and constitutively active (CA) RHOB was generated by PCR substituting glycine-14 with valine (G14V) in the GDP/GTP binding site^82,83^. We diluted 1 µg of plasmid DNA in 50 µL of Xfect Reaction Buffer (Takara Bio, # 631318), and then added 0.3 µL of Xfect Polymer to the diluted DNA. After incubation for 10 min at room temperature to form the transfection complex, the prepared transfection complex was gently added to each well. After 4 h incubation at 37 °C in a humidified incubator with 5% CO₂, we replaced the transfection medium with fresh growth medium and incubated to recover and express RHOB for 48 h.

For validation of RHOB expression, we added the substrate for Nluc (99-325-50, NanoLight Technology) to the culture medium according to the reagent’s instructions and measured the luminescence using a microplate reader (Nivo, Perkin Elmer) to compare the luminescence intensity between the experimental group (RHOB WT, RHOB CA, or RHOC WT plasmid) and controls (empty vector). After validation, we confirmed cell growth, phagocytic activity, actin polymerization, and cell adhesion, and investigated the effect of rhosin on RHOB overexpression-suppressed TM phagocytosis.

### β-galactosidase assays

To measure SA-β-galactosidase (senescent cell marker), we used the Cellular Senescence Detection Kit -SPiDER-βGal protocol (#SG02, DOJINDO), following the manufacturer’s instructions. Briefly, dobutamine and Dex at final concentrations of 0, 0.001, 0.01, 0.1, 1, and 10 µM were added to WT and RHOB KO iHTMCs 1 d after the medium was changed to serum-free TMCM. After 6 h, dobutamine-treated iHTMC were incubated with SPiDER-βGal working solution (1:1,000) at 37 °C for 30 min after washing twice with HBSS. iHTMCs were exposed to Dex for 96 h. After washing once with HBSS, fluorescence (Ex = 500 nm; Em = 540 nm) was measured using a Nivo spectrophotometer (PerkinElmer).

### Statistics and reproducibility

The results are shown as the mean ± standard error of the mean (SEM) of at least three independent experiments and five mice. Statistical comparisons were performed using GraphPad Prism 8 software (GraphPad Software Inc., San Diego, CA, USA). Paired or Student’s *t*-tests were used to compare two groups, one-way analysis of variance (ANOVA) with Dennett’s multiple comparison test for more than three groups, and two-way ANOVA with Sidak or Dennett’s multiple comparison test for more than three groups with two factors. Differences were considered statistically significant at *p* < 0.05.

## Data availability statement

Gene expression data in human AH outflow-related TM cells were deposited under the Gene Expression Omnibus accession number GSE146188^32^. Microarray data and RNA-seq data were deposited under the DDBJ accession number PRJDB20328 and PRJDB20330, respectively. All data are available from the corresponding author upon request.

## Acknowledgments

We are grateful to the Institute of Comprehensive Medical Research, Division of Animal Research Promotion Division (Aichi Medical University) for maintaining the mice, providing advanced research promotion for performing the F-actin fluorescence measurements, and to the Center for Advanced Technical and Educational Supports, Faculty of Agriculture, Kyushu University, for maintaining the mice, for the use of Nivo microplate reader, and for the expression analysis of proteins. We would like to thank Editage (www.editage.com) for English language editing. This study was supported by JSPS KAKENHI (Grant Number 19K09962), Takeda Science Foundation, Suzuken Memorial Foundation, Kato Memorial Bioscience Foundation, UBE Foundation, Yokoyama Foundation for Clinical Pharmacology, The Hori Sciences and Arts Foundation, the Toyoaki Scholarship Foundation (K.I.), and JST Fusion-Oriented Research for Disruptive Science and Technology (FOREST) (grant numbers JPMJFR215U) (A.O.).

## Author contributions

K.I. contributed to the conceptualization, methodology, investigation, writing of the original draft, and funding acquisition. T.T contributed to investigation. A.O., A.Y., and M.N. contributed to writing the review, editing and construct a series of plasmids. K.I., A.M., S,Y., and S.M. contributed to writing the review, editing.

Supplementary Information is available for this paper.

